# Prophylactic and therapeutic HBV vaccination by an HBs-expressing cytomegalovirus vector lacking an interferon antagonist

**DOI:** 10.1101/2020.01.29.924787

**Authors:** Hongming Huang, Meike Rückborn, Vu Thuy Khanh Le-Trilling, Dan Zhu, Shangqing Yang, Wenqing Zhou, Xuecheng Yang, Xuemei Feng, Yinping Lu, Mengji Lu, Ulf Dittmer, Dongliang Yang, Mirko Trilling, Jia Liu

## Abstract

Cytomegalovirus (CMV)-based vaccines show promising effects against chronic infections in non-human primates. Therefore, we examined the potential of HBV vaccines based on mouse CMV (MCMV) vectors expressing the small HBsAg. Immunological consequences of vaccine virus attenuation were addressed by either replacing the dispensable gene *m157* (‘MCMV-HBs’) or the gene *M27* (‘ΔM27-HBs’), the latter encodes a potent interferon antagonist targeting the transcription factor STAT2. *M27* was chosen, since human cytomegalovirus (HCMV) encodes an analogous gene product, which also induced proteasomal STAT2 degradation by exploiting Cullin RING ubiquitin ligases. Vaccinated mice were challenged with HBV through hydrodynamic injection. MCMV-HBs and ΔM27-HBs vaccination achieved accelerated HBV clearance in serum and liver as well as robust HBV-specific CD8+ T cell responses. When we explored the therapeutic potential of MCMV-based vaccines, especially the combination of ΔM27-HBs prime and DNA boost vaccination resulted in increased intrahepatic HBs-specific CD8+ T cell responses and HBV clearance in persistently infected mice. Our results demonstrated that vaccines based on a replication competent MCMV attenuated through the deletion of an interferon antagonist targeting STAT2 elicit robust anti-HBV immune responses and mediate HBV clearance in mice in prophylactic and therapeutic immunization regimes.

## INTRODUCTION

More than 2 billion individuals have been infected with hepatitis B virus (HBV) worldwide. The prophylactic HBV vaccine is a tremendous medical success, which saved countless lives. However, due to its inability to confer therapeutic protection, chronic HBV infections remain a major public health issue affecting approximately 250 million individuals (1). Persisting HBV predisposes to end-stage liver diseases, such as liver cirrhosis and hepatocellular carcinoma. According to the WHO, HBV is responsible for more than 850,000 deaths per year (https://www.who.int/news-room/fact-sheets/detail/hepatitis-b) (2). Two types of antiviral strategies are currently available for chronic hepatitis B (CHB): PEGylated interferon alpha 2 (PEG-IFNα2) and nucleot(s)ide analogues (NUC), such as Entecavir and Tenofovir. However, both are suboptimal. The treatment with PEG-IFNα2 is associated with significant side effects (e.g., flu-like symptoms, mood disorders, and depression), and long-term virus clearance is limited to approximately one third of treated patients (3). NUC medication selects for viral resistance mutations and is hampered by frequent episodes of rebounding viremia after cessation of antiviral therapy (4). Therefore, alternative strategies for the treatment of chronic HBV infection are urgently needed.

The clearance of HBV by the immune system relies on a potent and broad T cell immune response, which usually becomes dysregulated during chronic HBV infection (5–7). This exhaustion of HBV-specific T cells is believed to be one major reason for the inability of the host to eliminate the persisting pathogen. Aiming to enhance the patient’s own antiviral cellular immune response by therapeutic vaccination, is considered a promising strategy. During the last two decades, countless attempts have been made to establish an effective therapeutic vaccine against CHB (8). For example, existing protein-based prophylactic vaccines have been given to chronically infected patients in order to restore HBV-specific immunity. Unfortunately, it turned out to be unsuccessful (9, 10). An antigen-antibody (HBsAg-HBIG) immune complex termed YIC initially showed promising results in preclinical models and in phases IIA and IIB clinical trials (11, 12). However, the results of a phase III clinical trial enrolling 450 patients were disappointing (13). A number of DNA-based vaccine regimens have shown promising efficacy in pre-clinical animal models, but also failed to generate effective therapeutic responses in humans (8, 14). Therefore, replicating virus-based vectors, which stimulate a broad range of immune responses including T cell-mediated immunity, gained increasing attention in the field of therapeutic HBV vaccine development. In particular, vectors based on adenoviruses (15), modified vaccinia virus Ankara (MVA) (16), and recombinant vesicular stomatitis virus (VSV) (17) are under investigation for treatment of CHB.

Recently, cytomegalovirus (CMV)-based vectors emerged as exciting platform for vaccines against infectious diseases and cancer (18). Rhesus CMV (RhCMV)-based vaccines provided protective immunity to rhesus macaques against simian immunodeficiency virus (SIV), ebolavirus (19), and tuberculosis (20). In successfully RhCMV-vaccinated animals, even a CD8+ T cell depletion did not result in detectable SIV rebound, indicating very efficient virus control (21, 22). Vaccination with RhCMV vectors elicits robust and long-lasting cellular immune responses against pathogens mediated by effector-memory CD8+ T cells (23). Importantly, RhCMV-based vaccines demonstrated their efficacy even in the presence of preexisting immunity against CMV (24). These findings prompted us to evaluate the potential of CMV as platform for prophylactic and therapeutic vaccines against CHB in the established HBV hydrodynamic injection (HDI) mouse model. Our recent findings indicate that the interferon (IFN) antagonism of CMVs relies on viral proteins targeting the transcription factor STAT2 for proteasomal degradation by exploiting the adapter protein DDB1 and cellular Cullin RING ubiquitin ligases(25–27). Virus mutants lacking these immune evasins are replication competent but highly attenuated *in vivo*(28). The protein pM27 mediates STAT2 degradation in the MCMV context, while HCMV encodes the protein pUL145 which acts as a functional analog of pM27 (Le-Trilling & Becker *et al*., *provisionally accepted*). The proteins pM27 and pUL145 share a functionally relevant H-box motif present in viral and cellular DDB1 Cullin-associated factors (DCAF). Based on these findings, we reasoned that the attenuation of CMVs through the loss of STAT2 antagonists may represent a promising approach to establish novel platforms for the vaccination against viruses such as HBV.

## RESULTS

### Construction and characterization of a recombinant MCMV expressing the HBV small surface antigen

In order to evaluate the vaccine performance of CMV-based vaccine vectors in small animal models, we inserted an expression cassette comprising the coding sequence of the small HBsAg (sHBsAg; YP_009173871.1; corresponding to serotype ‘ayw’ HBV genotype D) under the control of the strong eukaryotic EF1 promoter into the mouse CMV (MCMV) genome by site-specific recombination. The expression cassette was introduced into an MCMV bacterial artificial chromosome (BAC) harboring a deletion of the coding sequences (CDS) of *m157* as schematically depicted in Fig. 1A and described in the Methods section. The gene *m157* is dispensable for MCMV replication *in vitro* and *in vivo* (29). To generate a live-attenuated vaccine virus, the expression cassette was also inserted into an MCMV mutant genome lacking the gene coding for the interferon antagonist pM27. The protein M27 interferes with interferon signaling by inducing ubiquitination and proteasomal degradation of STAT2 (26). *M27*-deficient MCMVs (‘ΔM27-Ctrl’) exhibit a slightly impaired (∼10-fold) replication in the absence of exogenous interferon treatment *in vitro*, but are severely attenuated *in vivo*(30, 31) in a STAT2-dependent manner (28). Correct mutagenesis was verified and HBsAg expression was confirmed by immunoblotting (Fig. 1B). As expected from previous work(32, 33) and the presence of an N-glycosylation site within the S domain of sHBsAg (Asn-146; sequence NcT), we observed two sHBsAg protein forms (Fig. 1B), most likely reflecting the previously described non-glycosylated p24 and the glycosylated gp27. The insertion of sHBsAg did not impair the replication of MCMV in cell culture (Fig. 1C). *In vivo*, we observed a trend towards reduced MCMV replication upon insertion of the sHBsAg expression cassette (Fig. 1D). As expected from our previous work, the deletion of *M27* resulted in a reduced replication *in vitro* (Fig. 1C) and a pronounced attenuation *in vivo* (Fig. 1D). Taken together, we generated two MCMV-based vectors expressing sHBsAg, one of which is severely attenuated *in vivo* due to the deletion of the MCMV-encoded STAT2 antagonist *M27*.

**Figure 1.**
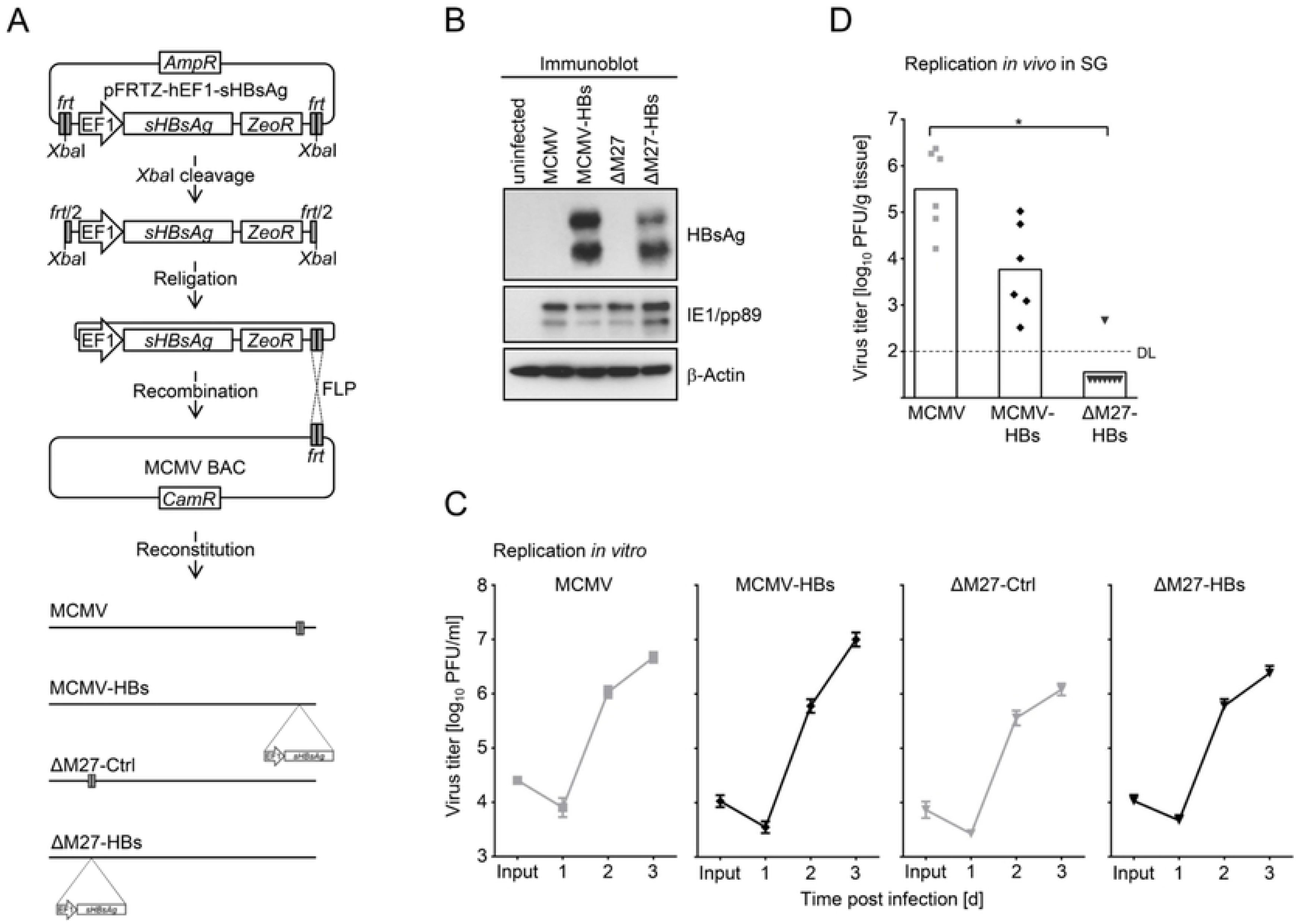
Generation and validation of HBsAg-expressing MCMV vectors. (A) Schematic overview of the cloning strategy for the generation of the HBsAg-expressing MCMV vectors. Please, see the M&M section for details. (B) HBsAg expression was confirmed by immunoblot. (C) The *in vitro* replication competence of MCMV vectors as well as corresponding parental viruses was determined by classic plaque titration. Mouse newborn cells were infected (0.05 PFU/cell) with indicated MCMVs. At indicated time points, cells and supernatants were frozen and stored at - 80°C until all samples were titrated simultaneously. Titrations were performed in triplicates. (D) The *in vivo* replication competence of the HBsAg-expressing MCMVs and wt-MCMV was compared in salivary glands at 21d post i. p. infection of C57BL/6 mice (n=6 mice per group).

### Vaccination with a recombinant MCMV expressing HBsAg protects mice against HBV challenge

We explored the protective effects of MCMV-HBs vaccination against an HBV challenge in the established HBV HDI mouse model. As schematically depicted in Fig. 2A, C57BL/6 mice were inoculated twice with MCMV-HBs, and then challenged with an HBV-expressing plasmid through HDI (see the Methods section for experimental details). Since the MCMV infection itself can negatively influence other infections (34) including HBV (35), the parental MCMV (‘MCMV’) devoid of the HBsAg expression cassette was included as negative control. While mice infected with empty MCMV exhibited continuous HBV viremia, the clearance of serum HBsAg, HBeAg, and HBV DNA was significantly accelerated in mice that received the MCMV-HBs vaccine (Fig. 2B). All MCMV-HBs-vaccinated mice became negative for serum HBsAg and HBeAg. Eighty percent of mice even became HBV DNA-negative at 9 days post HDI (dpi), while all control mice remained viremic for HBsAg, HBeAg, and HBV DNA (Fig. 2B and 2C). The MCMV-HBs-vaccinated mice also cleared HBsAg and HBcAg from the liver at 10 dpi, while the control mice expressed high levels of HBsAg and HBcAg, as evident by immune-histochemical staining (Fig. 2D and 2E).

**Figure 2.**
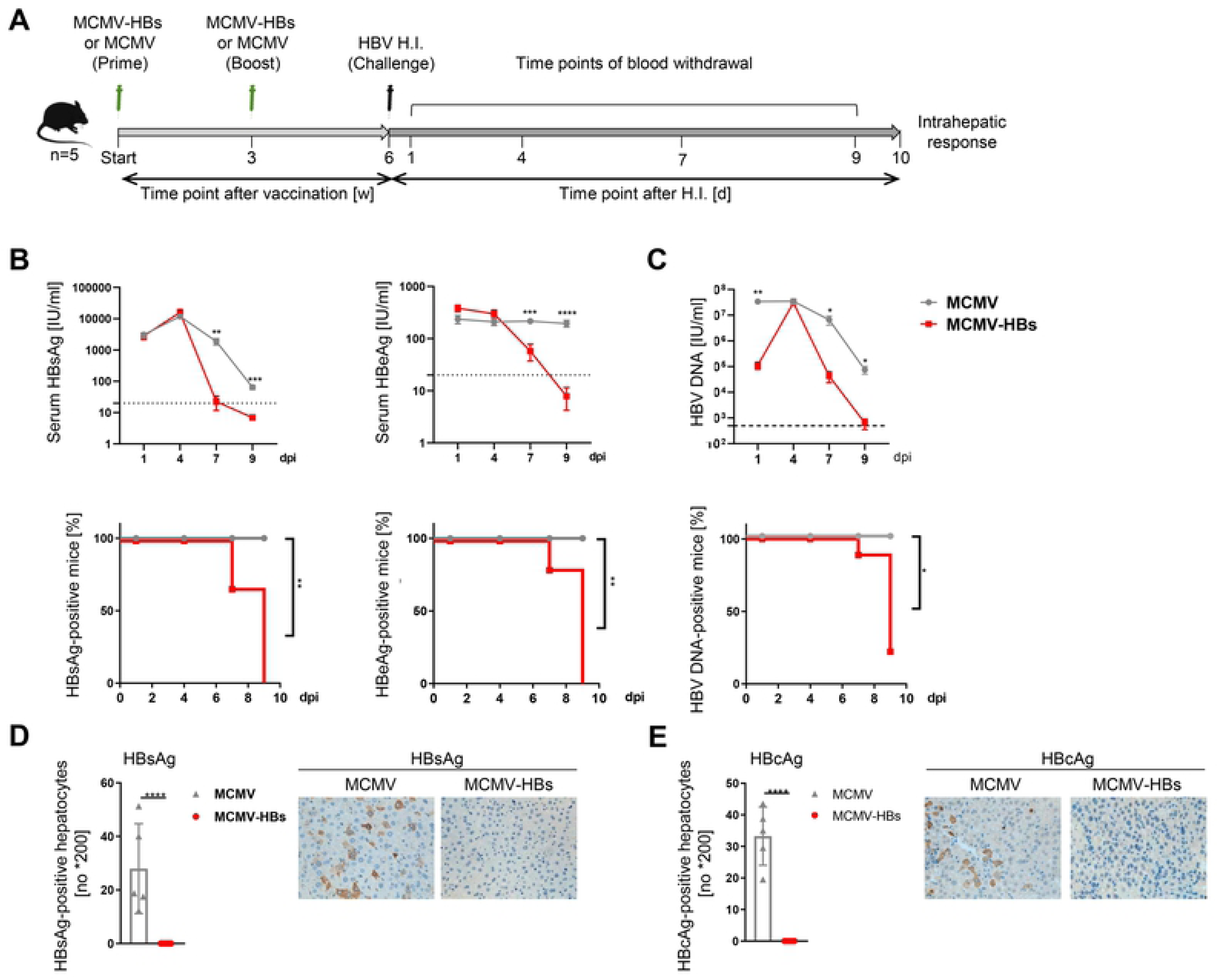
An HBsAg-expressing MCMV protects mice against HBV challenge. (A) Schematic overview of the experimental setup: C57BL/6J naïve mice were immunized intraperitoneally with 2*10^5 PFU of wt-MCMV or MCMV-HBs at day 0 and day 21 and were challenged with HBV plasmid pSM2 via hydrodynamic injection (HI) at day 42. The mice were sacrificed after HBV clearance in serum at 52dpi, 10 days after HI. (n=5). (B) The kinetics of serum HBsAg and HBeAg levels were monitored by ELISA at indicated time points. (C) The kinetics of serum HBV DNA levels were monitored by real-time PCR at indicated time points. (D) Immunohistochemical staining of HBsAg in the livers of wt-MCMV and MCMV-HBs immunized mice. Left: calculation of the average numbers of HBsAg-positive hepatocytes per field of vision. Right: representative staining of a liver section (original magnification: 200×) (E) Immunohistochemical staining of HBcAg in the livers of wt-MCMV- and MCMV-HBs-immunized mice. Left: calculation of the average numbers of HBcAg-positive hepatocytes per field of vision. Right: representative staining of a liver section (original magnification: 200×) Data are depicted as arithmetic means ± SEM, and experiments were repeated three times (n=5-6 mice per group). The statistical analysis was performed by Log-rank test (B-C) or unpaired t test (D-E). dpi, days post infection

### Vaccination with a recombinant MCMV expressing HBsAg raises potent CD8+ T cell responses

Next, we examined the impact of MCMV-HBs vaccination on immune responses. No anti-HBsAg antibodies were detectable in serum until day 7 after HDI for MCMV-HBs vaccination and day 9 after HDI for wt-MCMV vaccination (data not shown). However, since the HBV challenge is administered in form of a DNA molecule in the HDI mouse model, we reasoned that protection cannot be explained by neutralizing antibodies. Therefore, we focused our attention on T cell responses. We addressed intrahepatic T cell infiltration, activation, and HBV-specific CD8 T cell immune responses. In terms of frequency and absolute numbers, MCMV-HBs-vaccinated mice showed significantly higher CD8+ T cell numbers in the liver, while CD4+ T cells remained largely unaffected (Fig. 3A). A phenotypic analysis revealed that PD-1, but not CD43, expression on CD8+ T cells in the liver of MCMV-HBs-vaccinated mice was significantly increased compared to control mice (Fig. 3B and 3C). Since PD-1 is an activation marker for CD8+ T cells during acute HBV infection (36), this result suggests that MCMV-HBs vaccination enhanced the CD8+ T cell activation in the liver of HBV-challenged mice. In the liver, the MCMV-HBs-vaccinated mice showed a significant increase in absolute numbers and frequencies of CD8+ T cells specific for HBsAg, but not HBcAg, compared to control mice, as indicated by cytometry using Env190- and Core93-dimers, respectively (Fig. 3D and 3E). Accordingly, significantly higher percentages and absolute numbers of CD8 T cells in the livers of MCMV-HBs-vaccinated mice produced IFNγ or IL-2 in response to stimulation with the HBsAg epitope peptide (Env190), but not HBcAg epitope peptide (Core93) (Fig. 3F and 3G), suggesting that MCMV-HBs specifically enhanced HBsAg-specific CD8+ T cell responses.

**Figure 3.**
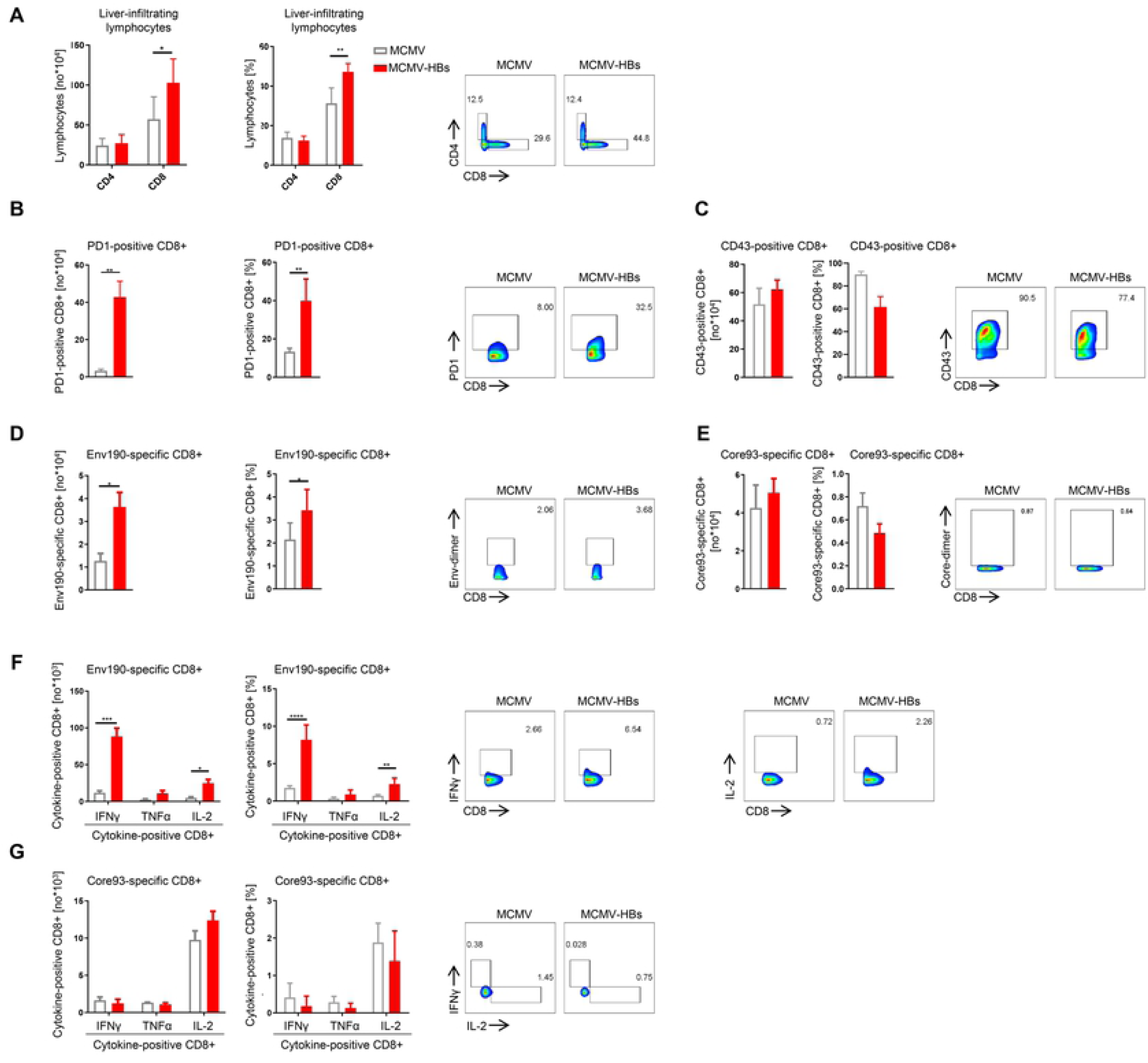
MCMV-HBs vaccination enhances the intrahepatic HBsAg-specific CD8 T cell responses. (A) Numbers, frequencies, and representative cytometry plots of CD4+ and CD8+ T cells among liver infiltrating lymphocytes in mice immunized with MCMV-HBs or MCMV. (B) Numbers, frequencies, and representative cytometry plots of PD1+ cells among CD8+ T cells in mice immunized with MCMV-HBs or wt-MCMV. (C) Numbers, frequencies, and representative cytometry plots of CD43+CD8+T cells among CD8+ T cells in mice immunized with MCMV-HBs or MCMV. (D) Numbers, frequencies, and representative cytometry plots of HBV Env190-specific CD8+ T cells. (E) Numbers, frequencies, and representative cytometry plots of HBV Core93-specific CD8+ T cells. (F) Numbers, frequencies, and representative cytometry plots of Env190-specific cytokine-positive CD8+ T cells after *in vitro* stimulation with Env190 peptide. (G) Numbers, frequencies, and representative cytometry plots of Core93-specific cytokine-positive CD8+ T cells after *in vitro* stimulation with Core93 peptide. Data were replicated in at least 2 independent experiments. Data are depicted as arithmetic mean ± SEM; *p <0.05, **p <0.01, ***p <0.001, ****p <0.0001; A unpaired t test was used to assess statistical significance; Core93, core93-100; Env190, env190-197;

### Vaccination with an HBsAg-expressing MCMV attenuated through the deletion of the interferon antagonist pM27 protects mice against HBV challenge

Even if cytomegaloviral vectors may be excellent vectors based on their ability to induce very strong T cell responses against foreign antigens, the use of replication competent HCMV-based vectors in humans is limited by their pathogenicity especially for immunocompromised individuals. One option to circumvent this safety issue is the use of live-attenuated CMV mutants(37), which may be achieved by different means. Therefore, we explored the potential of an HBsAg-expressing MCMV vector, which is replication-competent in cell culture, but highly attenuated *in vivo* due to the inability to counteract STAT2-dependent IFN signaling resulting from the lack of the gene product pM27 (28, 30, 31). C57BL/6 mice were inoculated twice with either empty ΔM27-MCMV (‘ΔM27-Ctrl’) or ΔM27-MCMV expressing HBsAg, for convenience denoted ‘ΔM27-HBs’ thereafter, and then challenged with an HBV-expressing plasmid through HDI. We also included PBS-treated mice as negative control to evaluate the protective effect of ΔM27-HBs (Fig. 4A). The clearance of serum HBsAg, HBeAg, and HBV DNA was significantly accelerated in mice vaccinated with ΔM27-HBs as compared to those which received PBS or the empty ΔM27-Ctrl (Fig. 4B and 4C). All ΔM27-HBs-vaccinated mice became serum HBsAg negative at 9 dpi, while the PBS-treated and ΔM27-Ctrl-infected mice remained 100% positive for serum HBsAg at this time point (Fig. 4B). Eighty percent of ΔM27-HBs-vaccinated mice also became HBeAg negative and 60% became serum HBV DNA negative at 9 dpi. In contrast, none of the mice from the PBS-treated or ΔM27-Ctrl-infected control groups cleared HBeAg or HBV DNA (Fig. 4C). The ΔM27-HBs-vaccinated mice were also the only animals that cleared HBsAg and HBcAg from the liver as evident by diminished immunohistochemical staining (Fig. 4D and 4E).

**Figure 4.**
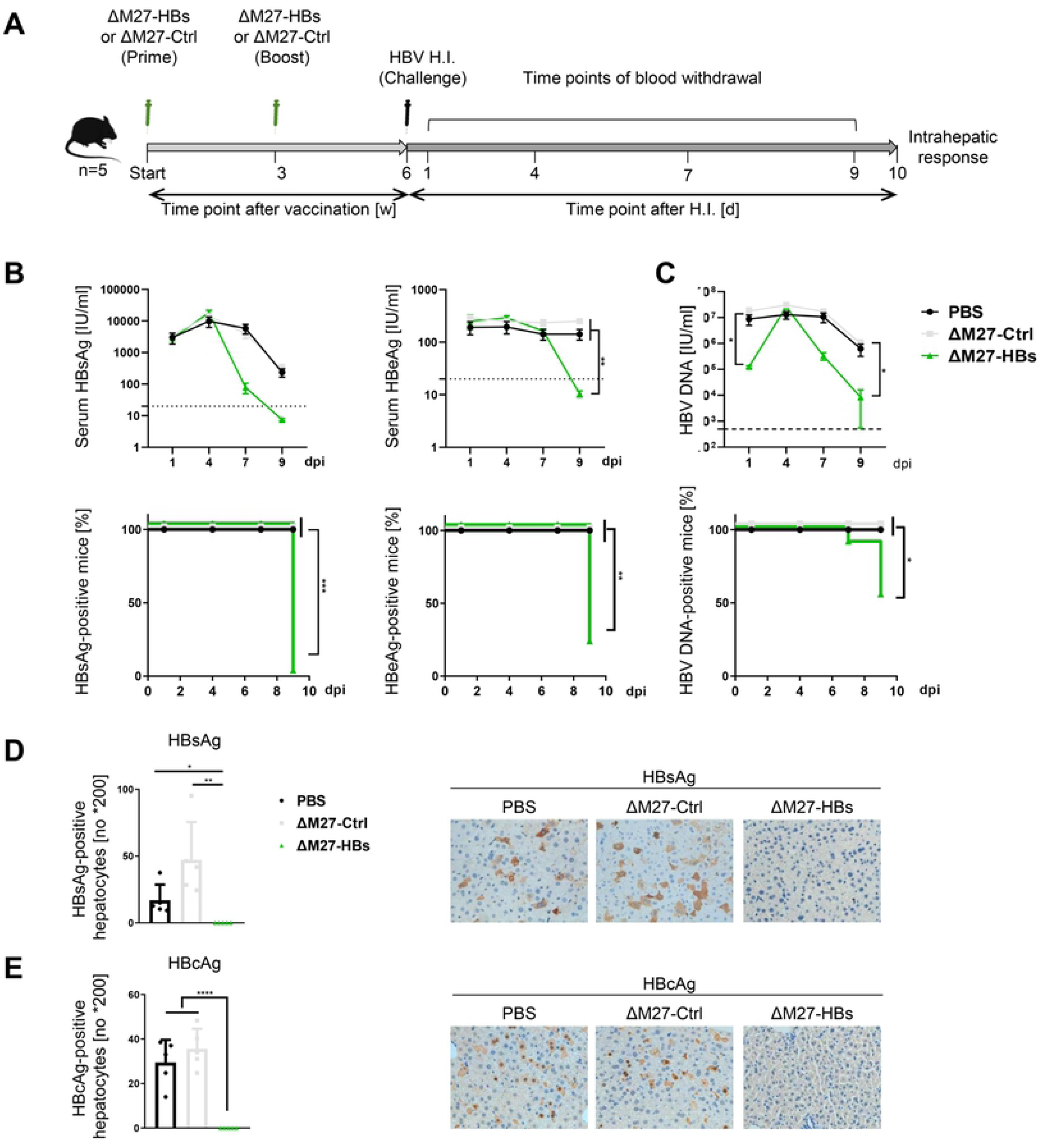
ΔM27-HBs vaccination protects mice against HBV challenge in mice. (A) Schematic overview of the experimental setup: C57BL/6J naïve mice were immunized intraperitoneally with 2*10^5 PFU of ΔM27-Ctrl, ΔM27-HBs or PBS in a volume of 100μL at day 0 and day 21, and were challenged with the HBV plasmid pSM2 via hydrodynamic injection (HI) at day 42. The mice were sacrificed after HBV clearance in serum at 52dpi, 10 days after HI. (n=5). (B) The kinetics of serum HBsAg and HBeAg levels were monitored by ELISA at indicated time points. (C) The kinetics of serum HBV DNA levels were monitored by real-time PCR at indicated time points. (D) Immunohistochemical staining of HBsAg in the livers of PBS-, ΔM27-Ctrl-, and ΔM27-HBs-immunized mice. Left: calculation of the average numbers of HBsAg-positive hepatocytes per field of vision. Right: representative staining of a liver section (original magnification: 200×) (E) Immunohistochemical staining of HBcAg in the livers of PBS-, ΔM27-Ctrl, and ΔM27-HBs-immunized mice. Left: calculation of the average numbers of HBcAg-positive hepatocytes per field of vision. Right: representative staining of a liver section (original magnification: 200×). Data are depicted as arithmetic means ± SEM, and experiments were repeated three times (n=5∼6 mice per group). The statistical analyses were performed by Log-rank test (B-C) or one-way ANOVA (D-E).

### Vaccination with an HBsAg-expressing MCMV attenuated through the deletion of the interferon antagonist pM27 induces potent CD8+ T cell responses

Compared to the PBS-treated control mice, the livers of ΔM27-HBs-vaccinated mice showed significantly higher percentages and absolute numbers of CD8 T cells, but not CD4+ cells (Fig. 5A). Interestingly, the ΔM27-Ctlr-infected mice also showed a significant increase in both percentages and absolute numbers of CD8+ T cells in the liver as compared to PBS-treated control mice (Fig. 5A, right panel), but the effect was more pronounced in ΔM27-HBs-vaccinated mice. Consistent with the above mentioned observation in the MCMV-HBs-vaccinated mice, PD-1 expression on CD8+ T cells in the liver of ΔM27-HBs-vaccinated mice was significantly increased compared to the PBS-treated or ΔM27-Ctrl-infected mice (Fig. 5B). Both ΔM27-Ctrl and ΔM27-HBs led to an increase in absolute numbers of CD43+ CD8+ T cells infiltrating the liver as compared to PBS-treated mice (Fig. 5C). These results suggest that the MCMV infection by itself results in enhanced infiltration of activated CD8+ T cells into the liver. Compared to the controls, the ΔM27-HBs-vaccinated mice showed a significant increase of both absolute numbers and frequencies of HBsAg-specific CD8+ T cells (Fig. 5D).

**Figure 5.**
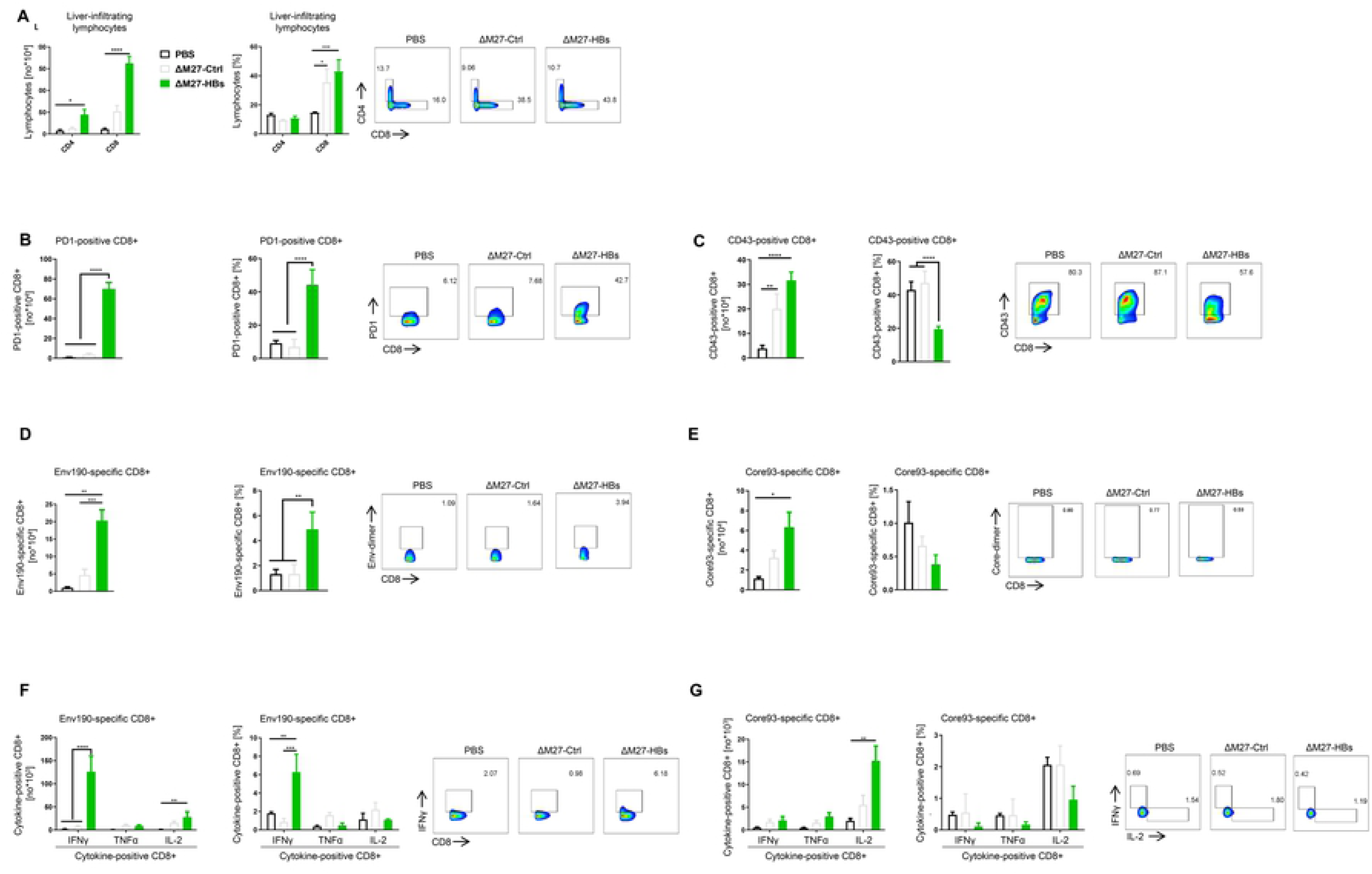
ΔM27-HBs vaccination enhances the intrahepatic anti-HBsAg CD8 T cell response. (A) Numbers, frequencies, and representative cytometry plots of CD4+ and CD8+ T cells among liver infiltrating lymphocytes in mice immunized with PBS, ΔM27-Ctrl, or ΔM27-HBs. (B) Numbers, frequencies, and representative cytometry plots of PD1+ cells among CD8+ T cells in mice immunized with PBS, ΔM27-Ctrl or ΔM27-HBs. (C) Numbers, frequencies, and representative cytometry plots of CD43+ cells among CD8+ T cells in mice immunized with PBS, ΔM27-Ctrl or ΔM27-HBs. (D) Numbers, frequencies, and representative cytometry plots of HBV Env190-specific CD8+ T cells. (E) Numbers, frequencies, and representative cytometry plots of HBV Core93-specific CD8+ T cells. (F) Numbers, frequencies, and representative cytometry plots of Env190-specific cytokine-positive CD8+ T cells after *in vitro* stimulation with Env190 peptide. (G) Numbers, frequencies, and representative cytometry plots of Core93-specific cytokine-positive CD8+ T cells after *in vitro* stimulation with Core93 peptide. Data were replicated in at least 2 independent experiments. Data are depicted as arithmetic mean ± SEM; *p <0.05, **p <0.01, ***p <0.001, ****p <0.0001; Statistical significance was assessed using the one-way ANOVA test; Core93, core93-100; Env190, env190-197;

The Core-specific CD8+ T cells were significantly increased in numbers but not in percentages in the liver of ΔM27-HBs vaccinated mice ten days after the challenge (Fig. 5E). When C57BL/6 mice were inoculated only once with MCMV-HBs or ΔM27-HBs, mice were also protected against a challenge with an HBV-expressing plasmid applied by HDI 3 weeks later (data not shown). Like in case of the prime-boost vaccination regime, single round vaccination with MCMV-HBs or ΔM27-HBs induced enhanced intrahepatic anti-HBV CD8+ T cell responses *in vivo* (data not shown).

Significantly higher percentages and absolute numbers of CD8+ T cells in the livers of ΔM27-HBs-vaccinated mice produced IFNγ in response to stimulation with the Env190 peptide than stimulated cells from the control groups (Fig. 5F). Additionally, the absolute numbers of IL-2-producing CD8+ T cells in the liver of ΔM27-HBs-vaccinated mice were also significantly increased in response to both Env190 and Core93 peptide stimulations (Fig. 5G).

Taken together, these results demonstrate that an HBsAg vaccination based on an MCMV vector incapable to interfere with STAT2-dependent IFN signaling enhances intrahepatic anti-HBV CD8 T cell responses *in vivo* and prophylactically protects mice against HBV challenges.

### MCMV-based HBsAg vaccination accelerates HBV clearance and enhances intrahepatic anti-HBV CD8 T cell responses in HBV persistent mice

After these promising results in prophylactic vaccination regimens, we examined the therapeutic capacity of MCMV-based HBsAg vaccination for the treatment of a previously established persistent HBV infection. The efficacy of MCMV-based HBV vaccine vectors was investigated by applying the established pAAV/HBV1.2 HDI mouse model, which mimics persistent HBV infections in humans (38). C57BL/6 mice were hydrodynamically injected with pAAV/HBV1.2. We chose a strategy, which combines an MCMV-based priming with a DNA-based booster immunization. The experimental setup is depicted in Fig. 6A. Treatment with MCMV-HBs prime and DNA boost resulted in a substantial reduction in HBsAg and HBV DNA levels in the serum (Fig. 6B and 6C). Eighty percent of MCMV-HBs-vaccinated mice became serum HBsAg negative and 60% became HBV DNA negative at 42 dpi. In contrast, only 20% of the wt-MCMV-infected control mice became serum HBsAg negative and none cleared the HBV DNA viremia (Fig. 6B and 6C). The MCMV-HBs-vaccinated mice also cleared HBsAg and HBcAg from the liver at 43 dpi, while the wt-MCMV-infected control mice still harbored high levels of HBsAg and HBcAg in the liver (Fig. 6D and 6E).

**Figure 6.**
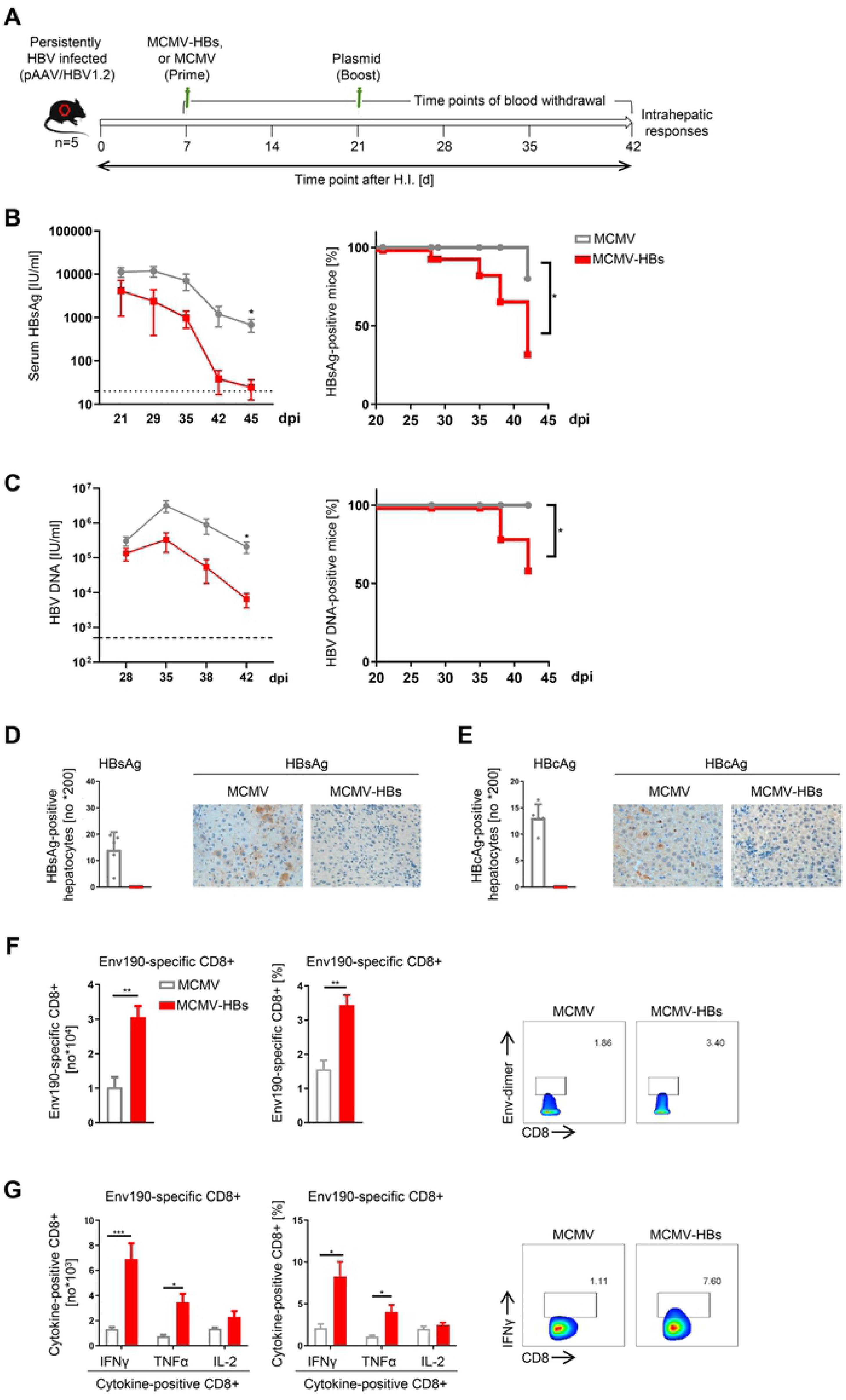
MCMV-HBs vaccination accelerates HBV clearance and enhances the intrahepatic anti-HBV CD8 T cell response in HBV persistent mice. (A) Schematic overview of the experimental setup: C57BL/6J naïve mice were hydrodynamically injected with pAAV-HBV1.2 (day 0) and immunized (primed) one week later (day 7) by intraperitoneal injection with MCMV-HBs or wt-MCMV and subsequently boosted with HBV plasmid pSM2 at week 4 (day 28). The mice were sacrificed after HBV clearance in serum at week 6, 42 days after HI. (n=5). (B) The kinetics of serum HBsAg levels were monitored by ELISA at indicated time points. (C) The kinetics of serum HBV DNA levels were monitored by real-time PCR at indicated time points. (D) Immunohistochemical staining of HBsAg in the livers of wt-MCMV- and MCMV-HBs-immunized mice. Left: calculation of the average numbers of HBsAg-positive hepatocytes per field of vision. Right: representative staining of a liver section (original magnification: 200×). (E) Immunohistochemical staining of HBcAg in the livers of MCMV- and MCMV-HBs-immunized mice. Left: calculation of the average numbers of HBcAg-positive hepatocytes per field of vision. Right: representative staining of a liver section (original magnification: 200×). (F) Numbers, frequencies, and representative cytometry plots of HBV Env190-specific CD8+ T cells. (G) Numbers, frequencies, and representative cytometry plots of Env190-specific cytokine-positive CD8+ T cells after *in vitro* stimulation with Env190 peptide. Data are depicted as arithmetic mean ± SEM, and experiments were repeated three times (n=5-6 mice per group). Statistical analyses were performed by Log-rank test (B-C) or unpaired t test (D-F).

Next, we examined the impact of MCMV-HBs vaccination on the intrahepatic anti-HBV CTL response. The percentages and absolute numbers of HBsAg-specific CD8 T cells were significantly increased in the liver of MCMV-HBs-vaccinated mice as compared to mice, which received the empty wt-MCMV (Fig. 6F). Moreover, significantly higher percentages and absolute numbers of liver CD8+ T cells of MCMV-HBs-vaccinated mice were capable of producing IFNγ and TNFα in response to Env190 peptide stimulation than cells derived from control treated mice (Fig. 6G). The same experimental setup was also applied to explore the therapeutic effect of ΔM27-HBs for the treatment of already-established persistent HBV replication (Fig. 7A). Interestingly, the attenuated ΔM27-HBs demonstrated even superior effects on accelerating HBV clearance than the more virulent and higher titer replicating MCMV-HBs in the HBV persistent pAAV/HBV1.2 HDI mouse model. All ΔM27-HBs-vaccinated mice became serum HBsAg and HBV DNA negative at 42 dpi, while the ΔM27-Ctlr-infected and PBS-treated mice remained 100% positive for serum HBsAg and HBV DNA at the time point (Fig. 7B and 7C). The ΔM27-HBs-vaccinated mice also cleared HBsAg and HBcAg from the liver at 43 dpi, while mice from the three control groups exhibited high levels of HBsAg and HBcAg in liver sections (Fig. 7D and 7E). Accordingly, ΔM27-HBs vaccination resulted in significantly increased infiltration of HBsAg-specific CD8+ T cells in the liver as compared to PBS treatment and ΔM27-Ctrl infection (Fig. 7F). Significantly higher percentages and absolute numbers of liver CD8+ T cells of ΔM27-HBs-vaccinated mice were capable of producing IFNγ in response to Env190 peptide stimulation than cells of control mice (Fig. 7G).

**Figure 7.**
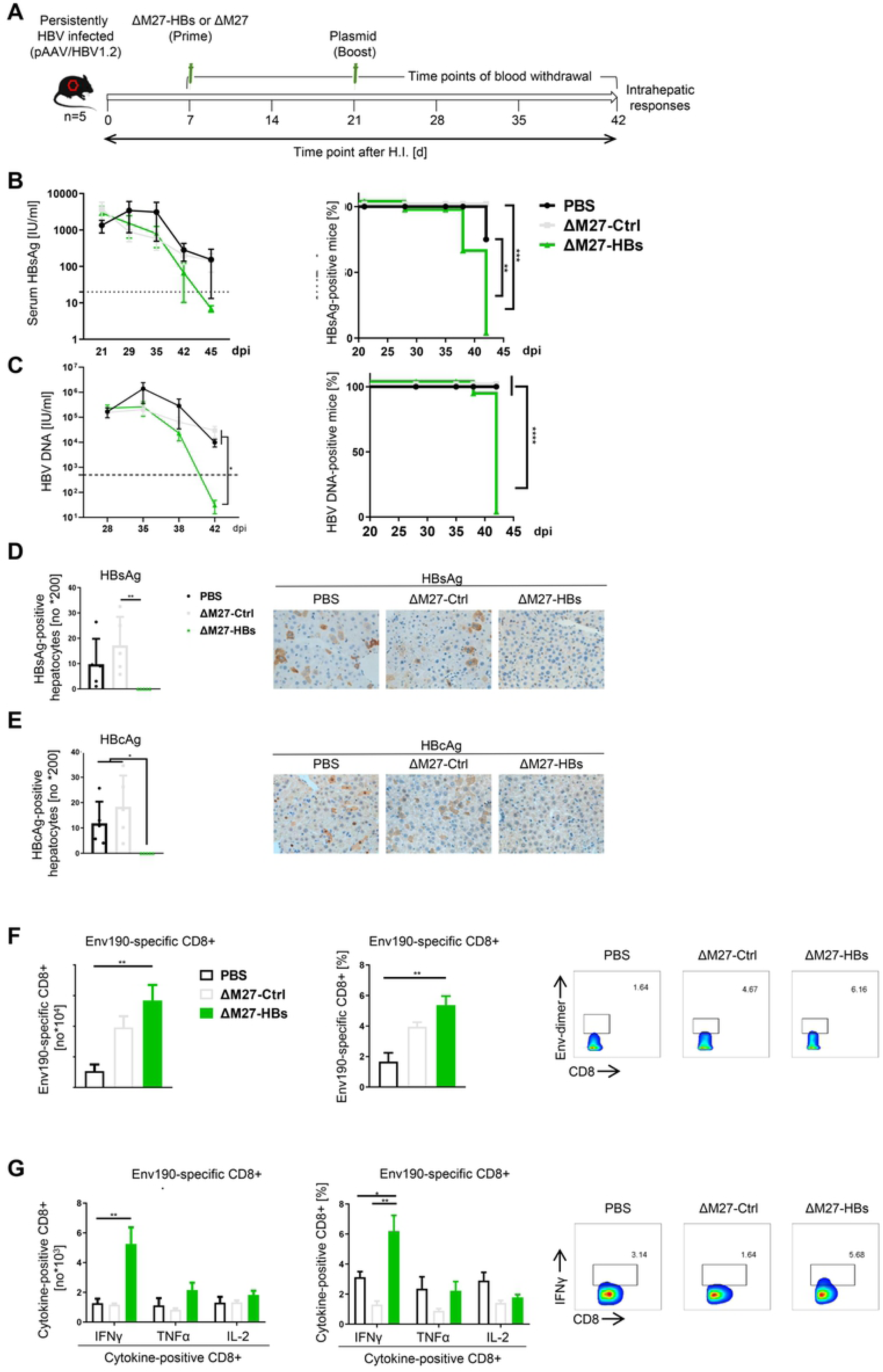
ΔM27-HBs vaccination accelerates HBV clearance and enhances the intrahepatic anti-HBV CD8+ T cell responses in HBV persistent mice. (A) C57BL/6J naïve mice were hydrodynamically injected with pAAV-HBV1.2 (day 0) and immunized (primed) one week later (day 7) intraperitoneally with 2*10^5 PFU of ΔM27-Ctrl, ΔM27-HBs, PBS in a volume of 100μL, and subsequently boosted with HBV plasmid pSM2 at week 4 (day 28). Mice were sacrificed after HBV clearance in serum at week 6, 42 days after HI. (n=5). (B) The kinetics of serum HBsAg levels were monitored by ELISA at indicated time points. (C) The kinetics of serum HBV DNA levels were monitored by real-time PCR at indicated time points. (D) Immunohistochemical staining of HBsAg in the livers of PBS-, ΔM27-Ctrl - or ΔM27-HBs-immunized mice. Left: calculation of the average numbers of HBsAg-positive hepatocytes per field of vision. Right: representative staining of a liver section (original magnification: 200×). (E) Immunohistochemical staining of HBcAg in the livers of PBS-, ΔM27-Ctrl - or ΔM27-HBs-immunized mice. Left: calculation of the average numbers of HBcAg-positive hepatocytes per field of vision. Right: representative staining of a liver section (original magnification: 200×). (F) Numbers, frequencies, and representative cytometry plots of HBV Env190-specific CD8+ T cells. (G) Numbers, frequencies, and representative cytometry plots of Env190-specific cytokine-positive CD8+ T cells after *in vitro* stimulation with Env190 peptide. Data are depicted as arithmetic mean ± SEM, and experiments were repeated three times (n=5-6 mice per group). Statistical analyses were performed by Log-rank test (B-C) or one-way ANOVA (D-F).

Taken together, our results suggest that HBsAg expression by an MCMV-based vector attenuated through the loss of the STAT2-specific IFN antagonist pM27 combined with a plasmid-based booster immunization could overcome HBV-specific CD8+ T cell dysfunction and induce HBV clearance in mice that previously established a persistent infection.

## DISCUSSION

HCMV is a betaherpesvirus that usually causes subclinical infections in healthy adults. However, HCMV elicits and sustains extraordinarily high numbers of antigen-specific T cells and is emerging as an exciting platform for vaccines against infectious diseases and cancers. To our knowledge, CMV-based vaccines against HBV have not been reported so far. Colleagues previously constructed an HBsAg-expressing MCMV (39), but seemingly did not approach the immunogenicity or vaccine potential. In the current study, we constructed two different recombinant MCMV vectors expressing HBsAg and explored their ability to induce HBV-specific CD8+ T cell responses and to inhibit HBV replication in HBV HDI mouse models. Our findings reveal that vaccination with the virulent MCMV-HBs and the attenuated ΔM27-HBs provided protection to mice against HBV challenges. The vaccinated mice quickly cleared HBV antigens and DNA from the serum and the liver, and generated robust intrahepatic HBsAg-specific CD8 T cell responses. Importantly, MCMV prime DNA boost vaccination strategies based on both recombinant MCMVs could even break HBsAg tolerance, elicit intrahepatic anti-HBV CD8+ T cell responses and lead to viral clearance in the serum and liver in persistently HBV-infected mice, suggesting that CMV-based vaccines possess therapeutic potential for treatment of chronic HBV infection in patients. In this respect, HCMV mutants lacking the pM27 analogous STAT2 antagonist pUL145 may constitute interesting candidates for live-attenuated vaccine platforms.

Prophylactic HBV vaccines that are currently in use rely on the induction of humoral immune responses against HBsAg, and thus neutralize infectious HBV particles before viral entry into hepatocytes. While antibodies are indispensable for prophylactic vaccination against HBV, this immune mechanism alone is most likely insufficient to eliminate HBV-infected cells in case vaccines are applied therapeutically, as indicated by the failures of approved prophylactic vaccines in therapeutic vaccination attempts. This assumption is consistent with our finding that the prophylactic HBV vaccine Engerix-B showed no protective effects in the acute HBV HDI mouse model (data not shown). However, this model is particularly biased against neutralizing antibodies since they are bypassed by unpackaged HBV genomes during direct transfection into hepatocytes through HDI. In case of a previously established infection, a potent cellular immune response mainly based on CD8+ T cells is essential for the elimination of infected cells (40). CMV infection causes an extraordinary strong T cell response. It has been estimated that CMV-specific T cell populations comprise an average 10% of the memory CD8+ T lymphocytes of infected individuals (41). In some cases, even up to 50% of all CD8+ T cells recognize a single epitope derived from the HCMV antigen pp65/pUL83 (42). Certain CMV-induced CD8+ T cell responses even show a so-called memory inflation, which means that their frequency continuously increases over time (43, 44). These inflationary CD8+ T cells are maintained as effector or effector-memory T cells, retaining the ability to produce inflammatory cytokines and to kill target cells (45, 46). In line with these reports, our findings demonstrate that vaccination with HBsAg encoded by MCMV vectors induced significantly increased numbers of activated HBsAg-specific CD8+ T cells infiltrating the livers of HBV-infected mice. These cells were capable of producing antiviral cytokines and led to accelerated HBV clearance. Interestingly, control MCMVs also induced increased infiltration of activated total, but not HBV-specific, CD8+ T cells in the liver. Since MCMV replicates efficiently in primary hepatocytes *in vitro* (see e.g., (47)) and the liver *in vivo* (see e.g., (28)), these results most likely reflect the effector nature of CMV in activating T cells and eliciting robust T cell responses against endogenous antigens. Such MCMV-specific T cell responses may explain why MCMV-HBs increases the absolute number of core-specific CD8+ T cells, but not the frequency (Fig. 5E). In addition, CMV possesses other features favorable for a vaccine vector. Firstly, HCMV can superinfect pre-immune hosts (23, 48). Thus, CMV-based vaccines may be applicable regardless of preexisting immune responses raised by naturally acquired CMVs, which would be favorable due to the high sero-prevalence of HCMV in countries with high incidences of CHB. Secondly, the large genome of CMVs allows CMV to take up large segments of foreign DNA at several different genomic loci. Another advantage might be the prolonged period of productive CMV replication, which can last several month in case of MCMV in mice (49), resulting in continuous antigen expression. The ability of herpesviruses to reactivate locally and spontaneously (50), leading to sequential rounds of immune stimulation, might further enhance potent immunogenicity.

Although most primary and recurrent HCMV infections progress subclinically in healthy adults, severe cases do occur (51). Additionally, HCMV causes morbidity and mortality in immune-immature, immune-compromised and immune-senescent individuals. Therefore, it is hard to envisage that authorities such as the FDA would approve vaccines based on virulent HCMVs. Thus, it will be necessary to increase the safety by attenuation. One appealing strategy for the generation of immunogenic and safe vaccines is the use of mutants lacking immune antagonists. Such viruses are by definition attenuated and therefore less likely to cause disease. Due to the inability to counteract a given aspect of immunity, such vectors may even induce superior immune responses. Here, we evaluated the effect of the lack the IFN antagonist pM27. ΔM27-MCMV is replication competent *in vitro* in the absence of IFN treatment but highly attenuated *in vivo* (28, 30, 31). Interestingly, the vaccination with the attenuated ΔM27-HBs showed superior protection compared to MCMV-HBs in terms of breaking the HBV tolerance in HBV persistent mice. Thus, the residual and temporary replication of the attenuated ΔM27-HBs vaccine generated sufficient numbers of HBsAg-specific CD8+ T cells to protect mice from the HBV challenge. Interferons stimulate the expression of several proteins involved in antigen processing and MHC presentation (see e.g., (52)). In conjunction with TCR activation and co-stimulatory molecules, IFNs can also serve as third signal for T cell activation (53). Thus, it is tempting to speculate that the inability of the MCMV vector to counteract STAT2-dependent IFN signaling is beneficial in terms of raising HBsAg-specific CD8 responses.

Altogether, our data corroborate the concept that CMV-based vectors are promising candidates for the development of new therapeutic vaccines for CHB treatment. The results of our study warrant more detailed investigations of HCMV-based vaccines to induce protective CD8 T cell responses against chronic HBV infection in patients.

## METHODS

### Genetic engineering of MCMV vaccine vectors

MCMV vectors were constructed by bacterial artificial chromosome (BAC) technology as described (^54, 55^). The coding sequence of the sHBsAg was amplified by PCR using the following primers, which introduce a C-terminal HA tag and harbour flanking restriction sites: MR-*Nhe*I-sHBsAg-fw (5’-tat *gct agc* atg gag aac atc aca tca gga ttc c-3’) and MR-sHBsAgHA-*Xho*I-rev (5’-ata *ctc gag* tta agc gta atc tgg aac atc gta tgg gta aat gta tac cca aag aca aaa gaa aat tgg-3’). The PCR product was cloned into the insertion plasmid pFRTZ-hEF1. The resulting pFRTZ-hEF1-sHBsAgHA harbours the HA-tagged sHBsAg gene under the control of the human elongation factor-1α (hEF1) promoter. Correct cloning was verified by restriction fragment analysis and sequencing (data not shown). pFRTZ-hEF1-sHBsAg was cleaved by *Xba*I which cleaves the two *frt* sites flanking the expression cassette and the *Zeo*^R^ gene. This fragment was religated, giving rise to a circular DNA element harbouring one *frt* site, the *Zeo*^R^ gene, and hEF1-sHBsAg, but lacking the plasmid backbone. The element was introduced into the *MCK-2* repaired MCMV-BACs (56) either harbouring an *frt* site instead of the gene *m157* or *M27* (28) by FLP-mediated homologous recombination in *E. coli* DH10B using the FLP-expressing plasmid pCP20. Correct mutagenesis was confirmed by restriction digest analysis, Southern blot and PCR analysis (data not shown). MCMV mutants were reconstituted by transfection of BAC DNA into permissive mouse newborn cells (57).

### Western Blot Analysis

Western blotting was performed as described previously (58) using the following antibodies: α-β-actin, α-HA (both from Sigma-Aldrich) and α-pp89-IE1 (CROMA101, generously provided by Stipan Jonjić, Rijeka, Croatia). Proteins were visualized using peroxidase-coupled secondary antibodies and an enhanced chemiluminescence system (Cell Signaling Technology).

### Mouse newborn cells(MNC)

MNC were prepared as described previously (57). In all experiments, MNC were used in passage 3. Cell culture media and supplements were obtained from Gibco/Life technologies and HyClone (Logan, UT). MNCs were used to propagate the MCMV viruses.

### MCMV propagation, titration, and replication analysis

MCMV stocks were generated as described (59). *In vitro* infections and titrations were enhanced by centrifugation (900g for 30 min). For the *in vivo* MCMV replication analysis, mice were infected intraperitoneally (i. p.). Organs of infected mice were harvested, snap-frozen in liquid nitrogen, and stored at -80°C until titration was performed.

### Mice

Male, 6- to 8-week-old wild-type C57BL/6J mice and timed pregnant 10- to 20-week-old female mice were purchased from Hunan Slack King Laboratory Animal Co., Ltd. (Changsha, China) and housed under specific pathogen-free (SPF) conditions in the Animal Care Center of Tongji Medical College or obtained from Charles River or Harlan and housed in the animal facility of the Institute for Virology of the University Hospital Essen. Timed pregnant 10- to 20-week-old female mice were used for generation of mouse newborn cells (MNC). Mice were sacrificed on the indicated days after vaccination and livers were harvested for analysis by IHC and flow cytometry. Procedures in Essen were conducted in accordance with regulations of the European Union and with permission of local authorities (the Landesamt für Natur, Umwelt, und Verbraucherschutz) in North Rhine Westphalia, Germany (permit numbers 84-02.04.2014.A390 & 84-02.04.2013.A414).

### Intraperitoneal injection in mice

Intraperitoneal injection in mice was performed using 2*10^5^ tissue culture infective dose affecting 50% of wells (TCID50) 50/ml of recombinant virus in 200μL (Δm157-MCMV [‘MCMV’], Δm157-MCMV: EF1-HBsAg [‘MCMV-HBs’], ΔM27-MCMV [‘ΔM27-Ctrl’], or ΔM27-MCMV: EF1-HBsAg [‘ΔM27-HBs’]). Identical volume of phosphate buffered saline (PBS) served as control.

### Hydrodynamic injection in mice

Hydrodynamic injection was performed as described previously using the replicating HBV plasmids pSM2 (generously provided by Dr. Hans Will, Heinrich-Pette-Institute, Hamburg, Germany) or pAAV/HBV1.2 (generously provided by Professor Pei-Jer Chen, National Taiwan University College of Medicine, Taipei, Taiwan) to establish HBV replication in mice (60–62). In brief, male mice (6 to 8 weeks of age) were injected with 10μg pSM2 or AAV/HBV1.2 in a volume of normal saline solution equivalent to 0.1 ml/g of the mouse body weight through the tail vein within 8 seconds (63, 64).

### Detection of serological HBV markers

Sera were prepared from blood collected at the indicated time points from the retro-orbital sinus of mice. Serum levels of HBsAg and HBeAg were measured by the corresponding ELISA kits (Kehua, Shanghai, China) according to manufacturer’s instructions. HBV DNA copies were determined by a diagnostic kit (Sansure, Changsha, China) using a quantitative real-time PCR according to manufacturer’s instructions.

### Cell isolation

Preparation of single-cell suspensions of murine intrahepatic lymphocytes was performed as described previously (64, 65).

### Flow cytometry

Surface and intracellular staining for flow cytometry analysis were performed as described previously (66). The following antibodies were used for surface and intracellular staining: FITC-anti-CD3, APC-Cy7-anti-CD4, Pacific Blue-anti-CD8, FITC-anti-CD43, PE-anti-PD1, and FITC-anti-IgG1 (all BD Biosciences, USA). For intracellular cytokine staining, we used the following antibodies: APC-anti-IFNγ, PerCP-Cy5.5-anti-IL-2, or FITC-anti-TNFα (BD Biosciences, USA). For HBV-specific CD8+ T cell detection, soluble DimerX H-2Kb: Ig fusion protein technology (BD Biosciences, USA) was applied. In brief, cells were incubated with CD16/CD32 anti-mouse antibody (clone 2.4G2; BD Pharmingen) to block FcRs. After washing, dimer staining was performed by incubation of dimer (which has been loaded with HBV Cor93-100 or Env190-197 peptide) and cells for 1 hour at 4°C. The cells were washed and incubated with anti-IgG1 antibody (BD Biosciences, USA) for 30 minutes at 4°C. Data were acquired on a BD FACS Canto II flow cytometer. Cell debris and dead cells were excluded from the analysis based on scatter signals and Fixable Viability Dye eFluor 506 (eBioscience). Isolated murine intrahepatic lymphocytes were used for all assays, and approximately 20,000-40,000 T cells were acquired for each sample using a BD FACS Canto II flow cytometer. Data analysis was performed using FlowJo software V10.0.7 (Tree Star, Ashland, OR, USA).

### Statistical analysis

Statistical data were derived by using the GraphPad Prism software (GraphPad Software). Data were analyzed using Log-rank test, unpaired t test and one-way ANOVA. As described in each figure legend.

### Ethics statement

All animal procedures were approved by the Institutional Animal Care and Use Committee Tongji Medical College, Huazhong University of Science and Technology in accordance with the recommendations in the National Advisory Committee for Laboratory Animal Research (NACLAR) guidelines. IACUC Number 2019-S1016. The experiments were performed under isoflurane anesthesia, and all efforts were made to minimize suffering.

## Acknowledgments

We thank Professor Pei-Jer Chen for providing the pAAV/HBV1.2 plasmid. This work was supported by the National Natural Science Foundation of China (91742114, 91642118, and 81861138044 and 91742114), the National Scientific and Technological Major Project of China (2017ZX10202203), and the Sino-German Virtual Institute for Viral Immunology. MT received funding from the Deutsche Forschungsgemeinschaft (DFG) through RTG1949 (project 13), TR1208/1-1, and TR1208/2-1. MT also received support from the Kulturstiftung Essen.

